# Coordinating the *in vivo* processes for minicircle production - a novel approach to large scale manufacturing

**DOI:** 10.1101/300236

**Authors:** Peter Mayrhofer, Hana Jug, Aleš Štrancar, Alexandre Di Paolo, Laurent Jost, Alice Zimmermann, Heinz Redl

## Abstract

Safety as well as efficiency issues in connection with bacterial backbone sequences should be carefully considered when designing new DNA vaccines or non-viral gene therapy approaches. Bacterial backbone sequences like antibiotic resistance markers or regulatory bacterial elements constitute biological safety risks and reduce the overall efficiency of the DNA agent. To overcome these problems the minicircle technology has been developed. But, despite all the obvious advantages, minicircles have so far not replaced their problem laden conventional counterpart in gene transfer applications what can be contributed to efficiency issues in large scale manufacturing. In this article we describe the combined efforts of experts in the field of minicircle development, large scale biomanufacturing and downstream process development to provide a new approach. The Recombination Based Plasmid Separation (RBPS) Technology, which has already solved crucial problems associated with minicircle-DNA production, has been developed further for this purpose. A novel parental plasmid exploiting advanced *in vivo* process coordination for restriction and subsequent degradation of miniplasmid-DNA will be introduced. Furthermore we describe the scale-up of minicircle-DNA production by fermentation in combination with high performance downstream processes including purification by ion exchange and hydrophobic interaction chromatography on monolithic material.

## INTRODUCTION

Minicircles are DNA-molecules derived from bacterial plasmid-DNA which are optimized for clinical gene transfer applications. Their specific advantage over their conventional counterparts is that bacterial DNA sequences constituting a biological safety burden have been removed via a sophisticated *in vivo* recombination reaction (Bigger et al. 2001; Chen et al. 2003; Darquet et al. 1997; Darquet et al. 1999; Jechlinger et al. 2004; Kreiss et al. 1998; Mayrhofer et al. 2008). It is now the dawn of the minicircle age in the field of non-viral gene delivery. Many studies are ongoing or have been conducted demonstrating the advantages of these DNA-molecules in specific medical applications e.g in the field of angiogenesis, lung treatment, rheumatoid arthritis or in the treatment for myocardial infarction just to mention a few (Bandara et al. 2016; Liu et al. 2017; Munye et al. 2016; Park et al. 2017). Furthermore, minicircle-DNA has already arrived in the field of CRISPR-Cas9 based gene editing (Schmid-Burgk et al. 2016), it is used as vector for RNA interference since years (Hornstein et al. 2016; Lijkwan et al. 2014) and has shown advantages in gene transfer applications using transposons like sleeping beauty (Hudecek et al. 2017; Sharma et al. 2013). Although the first publications on minicircle-DNA appeared decades ago there is still research ongoing to improve delivery strategies (Gaspar et al. 2015b; Wan et al. 2013; Wang et al. 2014), purification (Gaspar et al. 2013) and production (Dong et al. 2013). Reviews discussing status and progress in the field of minicircle-DNA are available (Gaspar et al. 2015a; Mayrhofer and Iro 2011). The production of minicircle-DNA is still not a straight forward process. The insufficiency of the involved processes has prevented the broad application of these molecules in gene transfer applications already for several decades. Although various technologies have been described for minicircle production, up-scaling for industrial manufacturing is still work in progress.

Conventional plasmid vectors consist of a bacterial backbone and a transcription unit whereas the latter carries the target gene or sequence along with necessary regulatory elements and the backbone DNA consists of elements like an antibiotic resistance marker, an origin of replication or unmethylated CpG motives that clearly need to be removed for clinical applications (Mayrhofer and Iro 2011). To remove these unwanted elements without destroying the supercoiled structure of the plasmid-DNA, the minicircle technology has been developed (Bigger et al. 2001; Chen et al. 2003; Darquet et al. 1997; Darquet et al. 1999; Jechlinger et al. 2004; Kreiss et al. 1998; Mayrhofer et al. 2008). This technology generates a circular, supercoiled, minimal expression cassette by an *in vivo* site-specific recombination process which removes the aforementioned backbone elements. Although *in vitro* minicircle-DNA production has also been reported (Dong et al. 2013; Thibault et al. 2017) the most common approach involve site specific recombination *in vivo* as the basis for a fermentation driven large scale manufacturing process. In the course of this process the parental plasmid (the producer plasmid) is divided into a miniplasmid carrying the backbone sequences and a minicircle consisting of almost exclusively the desired expression cassette (minimal expression cassette). With regard to the recombination, basic parameters influencing the efficiency like the type of recombinase used, its regulation as well as the location of the corresponding expression cassette have been investigated to result in a scalable production process (for review see (Mayrhofer and Iro 2011)). Other optimisation approaches included the modification specific components like for example the *loxP* restriction sites to avoid adverse recombination events mediated by the Cre recombinase (Bigger et al. 2001). A more recent publication reported the improvement of the secondary structure of the 5’-UTR of the ParA mRNA to achieve better expression of the resolvase (Simcikova et al. 2016).

Following the site-specific recombination process the resulting mixture of plasmid species (minicircles, miniplasmides and to some extent unrecombined parental plasmid) must be separated, to isolate the desired minicircle-DNA. Different strategies have been developed for this purpose including affinity based chromatographic purification (Mayrhofer et al. 2008) and *in vivo* restriction (Chen et al. 2005; Kay et al. 2010). Both techniques have pros and cons. Although the concept of *in vivo* restriction is very promising, the results published so far do not demonstrate the suitability for large scale minicircle-DNA production. Chen et al (Chen et al. 2005) reported in 2005 that there would remain about 3 % contaminating parental plasmid after 240 min. of recombination and *in vivo* restriction. In a more recent publication from 2010 this value was corrected to up to 15 % residual parental plasmid but for the improved system described in this work fewer contamination of about 1.5 % were reported (Kay et al. 2010). One drawback of the *in vivo* restriction system established so far is the lack of temporal coordination of the involved processes. In an ideal system, the processes of recombination and degradation should be performed consecutively, i. e. restriction should only start when the recombination process is complete, to avoid the premature degradation of the parental plasmid. So far these processes were induced simultaneously as the enzymes needed for recombination and restriction were both under the control of the same promoter/operator unit. This parallel induction of the involved enzymes resulted in premature restriction of the parental plasmid (Chen et al. 2005; Kay et al. 2010).

Various measures were taken to minimize premature degradation of the parental plasmid including the introduction of additional copies of the enzyme driving recombination, shifting temperature and changing the pH of the culture medium during the production process to favour either the recombination process or the restriction process (Chen et al. 2005; Kay et al. 2010). With regard to scale up and large scale biomanufacturing of minicircle-DNA an alternative way to coordinate these processes *in vivo* is needed as such changes cannot be performed so easily as in lab scale experiments (e.g.one litre shaking flask cultures). This problem has been solved by the next generation RBPS parental plasmid described in this work which enables consecutive recombination and *in vivo* restriction.

For this work the novel RBPS parental plasmid has been used in a scalable 5 litre fed batch fermentation approach. The resulting biomass was used to develop a scalable downstream process for minicircle-DNA purification including pre-chromatographic lysis and clarification steps and chromatographic capture and polishing steps using monolithic ion exchange (IEC) and hydrophobic interaction (HIC) chromatography systems.

## MATERIALS AND METHODS

DNA manipulations: Preparation of plasmid DNA and isolation of DNA fragments was carried out using kits from QIAGEN (High Speed Plasmid Maxi Kit, QIAquick PCR Purification Kit and QIAquick Gel Extraction Kit) and PEQLAB (peqGOLD Plasmid Mini Prep Kit I). Transformation of *E. coli* cells was either performed as described previously (Sambrook et al. 1989) or according to the supplier’s instructions. Competent *E. coli* NEB10-beta and NEB 5-alpha F’Iq were purchased from New England Biolabs and transformation was performed according to the supplier’s instruction. Electrophoresis of DNA was performed as described previously (Sambrook et al. 1989). If not otherwise stated, restriction enzymes, DNA-modifying enzymes and nucleotides were obtained from New England Biolabs or Promega, polymerases used for PCR reactions were purchased from Fermentas (Pfu) or Stratagene (Herculase) and used as specified by the manufacturer. Oligonucleotides for PCR and cloning were purchased from Sigma-Aldrich. Sequence analysis was accomplished by AGOWA.

Preparation of Buffers: Buffers for cell lysis, clarification step and chromatography were prepared fresh with dH_2_O. Buffers for chromatography were additionally filtrated with PES 0.22um filter (TPP, Trasadingen, Switzerland). TRIS (Trizma base), CaCl_2_, KCH_3_COOH, NaCl, SDS, (NH_4_)_2_SO4 and NaOH were obtained from Merck (Darmstadt, Germany), EDTA (Na2x2H2O komplexal III) from Kemika (Zagreb, Croatia).

Chromatographic equipment: Chromatography was performed using a Knauer HPLC system (Knauer, Berlin, Germany) consisting of two preparative pumps (1000 ml), an interface and Knauer UV-VIS absorbance detector K-2500 with 2 mm optical cell. Some of the experiments were performed using an Äkta Purifier 10 system (GE Healthcare Life Sciences, Boston, USA) with 10 mm optical cell.

Bacterial strains: Cloning work was generally carried out in *E. coli* NEB10-beta or *E. coli* NEB 5-alpha F’Iq purchased from New England Biolabs. The *E. coli* B strain BL21 F– ompT gal dcm lon hsdSB(rB–mB–) [malB+]K-12(λS) used for minicircle production was purchased from Novagen (Madison, Wisconsin, US)

### Cloning work

Due to the high number of necessary cloning steps the material and methods section provides a summary of the total cloning work. The details of the single cloning steps are available as supplementary material to this publication. Figure S1 which is available in the supplementary material provides a graphical overview of the cloning steps.

### Cloning of endonuclease I-TevI

#### Cloning of the I-TevI open reading frame into plasmid pACYC177

A PCR generated DNA fragment containing the open reading frame of I-TevI was purchased from GenScript (New Jersey, USA). This fragment was amplified by PCR and subcloned into vector pACYC177.

#### Cloning of a functional I-TevI expression cassette

To clone a functional I-TevI expression cassette under the control of the promoter/operator system of the arabinose operon (araBAD), the open reading frame of I-TevI was cloned into plasmid pLacOsIivb described by Mayrhofer (Mayrhofer et al. 2008). A PCR fragment containing the open reading of I-TevI and a frame a Shine-Dalgarno sequence was generated using pACYC177ITevI as template and cloned into plasmid placOsIivb. The resulting plasmid placOsSDEITevivb contains an I-TevI expression cassette using the Shine Dalgarno sequence of the lysis gene E of the *E. coli* bacteriophage PhiX174 under the control of the araBAD promoter of the promoter/operator system of the arabinose operon fused to an *in vivo* biotinylation sequence as described by Mayrhofer (Mayrhofer et al. 2008). Expression of the biotinylated fusion protein was verified by western blotting using streptavidin-alkaline phosphatase conjugate for detection (data not shown).

### Cloning of RBPS *in vivo* restriction vectors

The RBPS-Technology *in vivo* restriction (RBPS-IVR) approach is based on the RBPS-System as published by Mayrhofer et al (Mayrhofer et al. 2008). Basically, the same elements for recombination and chromatographic purification were used to build plasmids for *in vivo* restriction. Additionally, a promoterless expression cassette of an endonuclease (I-TevI) derived from a T-series *E. coli* phage, as well as the corresponding restriction site, were integrated in the novel parental plasmid (pRBPS-IVR). The I-TevI restriction sites were integrated at positions that are located on the miniplasmid after recombination. Upon expression of the endonuclease, plasmids carrying these corresponding sites were linearized and subsequently further degraded by host exonucleases. The elements on the parental plasmid were arranged in a way that the endonuclease expression cassette was activated by the removal of the minicircle region from the parental plasmid via the recombination process. Expression was achieved by switching a promoter element in front of the I-TevI cassette through the ParA driven plasmid rearrangement. As a result of this rearrangement, a miniplasmid carrying a functional I-TevI expression cassette was generated enabling the expression of this enzyme and consequently the degradation of miniplasmids.

#### Cloning of plasmidpRBPS-IVR1

In the first attempt it was planned to use the already available arabinose promoter/operator system regulating the ParA expression in pRBPS7 to drive I-TevI expression after the recombination event. In order to achieve this it was necessary to reverse the orientation of the arabinose P/O system so that the araBAD promoter (P_BAD_) would be switched in front of the I-TevI cassette (see Figure 1 “From pRBPS7 to pRBPS-IVRT”).

**Fig. 1a.**
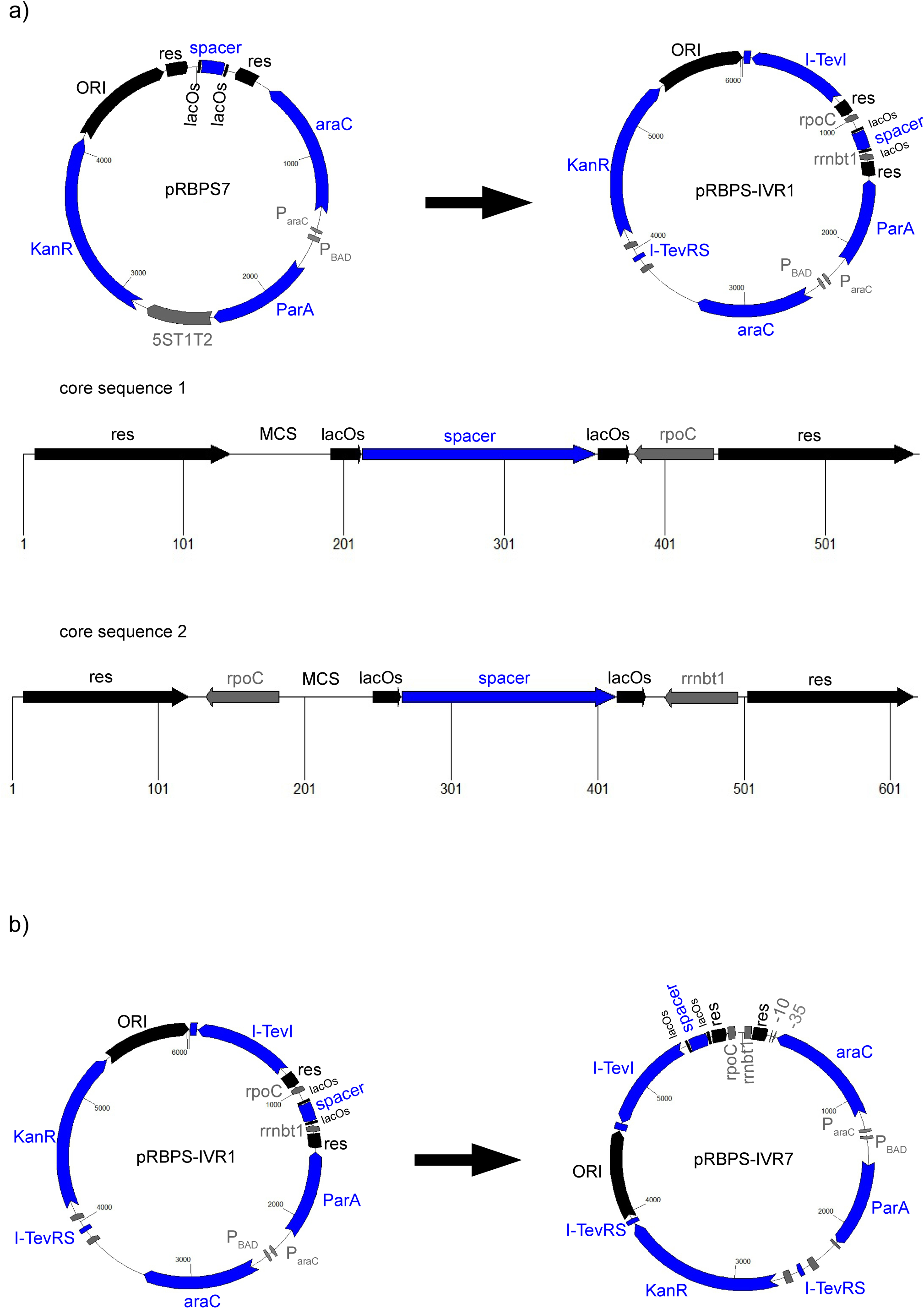
shows plasmid maps of pRBPS7 and pRBPS-IVR1. pRBPS7 served as the origin vector to generate the *in vivo* restriction plasmid pRBPS-IVR1. Core sequence 1 was integrated in plasmid pRBPS7_rrnbt1_rs to generate plasmid pRBPS18_rev_rs. Core sequence 2 was integrated in plasmid pRBS18_rev_rs_ITev to construct plasmid pRBPS-IVR1. **Fig. 1b** shows the plasmid maps of pRBPS-IVR1 and pRBPS-IVR7 Abbreviations: res, resolution site of the ParA resolvase system; lacOs, modified lactose operator site; spacer, spacer sequence between two direct repeats of the modified lactose operator sites lacOs; ParA, parA resolvase gene; ParaC, promoter of the araC gene; Pbad, promoter of the araBAD genes of the arabinose operon; araC, gene enconding the repressor of the arabinose operon; ITev-RS, ITev restriction site; Kan, aminoglycoside 3’-phosphotransferase expression cassette conferring resistance to kanamycin; ORI, MB1 origin of replication; ivb, *in vivo* biotinylation sequence; ITev, expression cassette of intron enconded endonuclease of bacteriophage T4; rpoC, transcription terminator rpoC; rrnbt1, transcription terminator rrnbt1; −10 −35, −35 and −10 region of the lacI^q^ promoter - an up-mutation of the constitutive promoter of the lacI gene of the lactose operon

To prepare for the rearrangement of the ParA expression cassette in plasmid pRBPS7 the spacer sequence between the lacOs sites was replaced by the terminator region of plasmid pBAD24 (Guzman et al. 1995) in the first step. This was achieved by inserting a PCR fragment containing this terminator region flanked by the lacOs sites into plasmid pRBPS1 (Mayrhofer et al. 2008) resulting in plasmid **pRBPS1/Ter** and subsequent subcloning of this fragment into pRBPS7 to generate plasmid **pRBPS7/Ter**. This vector was used as a precursor for the inversion of the ParA expression cassettes. To inverse the ParA expression cassette including its regulatory sequences a PCR fragment containing these elements was generated and inserted in reverse orientation into pRBPS7/Ter resulting in plasmid **pRBPS7_Ter_Rev**. Plasmid pRBPS7_ter_rev contains the full 426 bp terminator region of pBAD24 between the lacOs sites. Additionally, a plasmid harbouring a shorter terminator sequence was generated by substituting this large terminator region with the 51 bp sequence of terminator rrnbT1 (Brendel et al. 1986). The rrnbT1 sequence was generated by oligo annealing and cloned into plasmid pRBPS1 resulting in plasmid **pRBPS1_rrbt1**. Subsequently, the rrnbT1 sequence was subcloned into pRBPS7_Ter_Rev thereby generating plasmid **pRBPS7_rrnbt1_rev**.

To introduce the recognition site of endonuclease I-TevI into the vector system a dsDNA fragment containing the 39bp I-TevI site (Chu et al. 1991) was generated by oligo annealing. This fragment was cloned into plasmid pRBPS7_rrnbt1_rev resulting in vector **pRBPS7_rrbt1_rs** containing the araB promoter orientated counter clockwise, the rrbt1 terminator and a 39 bp restriction site of T-TevI.

In the next step a promoterless I-TevI expression cassette should have been subcloned from plasmid placOsSDEITevivb (described above) into one of the plasmids generated for this purpose (pRBPS7_Ter_Rev, pRBPS7_rrnbt1_rev or plasmids pRBPS pRBPSS7_rrnbT1_rs). However, this cloning step could not be successfully performed.

Further analysis revealed that the promoter sequences contained in the res sites (Eberl et al. 1994) show some activity leading to premature expression of the I-TevI (data not shown). Obviously, this background expression hampered the cloning of the promoterless I-TevI expression cassette. Therefore, these promoter elements were removed by exchanging single base pairs in the respective regions. From the *StyI/NdeI* fragment of the resolution site described in (Eberl et al. 1994), which is used for RBPS-Technology the −35 bp region of the parDE promoter region has been completely removed. The −10 and −35 bp regions of the parCBA promoter sequence were eliminated by single nucleotide exchanges (A -> G in the −10 bp and two vicinal A -> G in the −35 bp region). An appropriate sequence containing the modified res sites and the restriction sites *Not*I and *BstAPI* for the integration into plasmid pRBPS7_rrnbT1_rs was designed and purchased from GeneArt (Regensburg, Germany). This sequence contained the two altered resolution sites flanking the transcription terminator rpoC (Brendel et al. 1986), a multiple cloning site and the lacOs – spacer – lacOs sequence of pRBPS7 (see Figure 1a, core sequence 1). This sequence was inserted into restriction sites *Not*I and *BstAP*I of plasmid pRBPS7_rmbt1_rs resulting in vector **pRBPS18_rev_rs** containing the modified resolution sites and the transcription terminator rpoC.

To clone the promoterless I-TevI expression cassette into plasmid pRBPS18_rev_rs an appropriate fragment was subcloned from vector placOsSDEITevivb. The resulting vector **pRBPS18_rev_rs_ITev** contained the I-TevI expression cassette orientated counter clockwise.

In the next step a reporter cassette was to be inserted into the multiple cloning site of the vector pRBPS18_rev_rs_ITev which contained all necessary elements of the *in vivo* restriction system to test the system set-up. Despite all attempts, it was not possible to integrate an appropriate expression cassette into the multiple cloning site of pRBPS18_rev_rs.

Therefore, modifications were necessary to enable the insertion of an expression cassette into the parental plasmid. A second transcription terminator was inserted into pRBPS18_rev_rs_ITev to prevent unintended transcription of ITevI from potential cryptical promoters within the sequence of the insert. The functional elements were arranged as follows: res site, rpoC, MCS, lacOs-spacer-lacOs, rrnbT1, res site (see Figure 1a, core sequence 2). An appropriate sequence containing the described functional elements was designed and purchased from GeneArt (Regensburg, Germany). This sequence was introduced into plasmid pRBPS18_rev_rs_ITev. The resulting vector **pRBPS-IVR1** contained the elements as described above including a second transcription terminator. Despite the new set-up, the insertion of an expression cassette into pRBPS-IVR1 to produce the corresponding minicircle was not possible.

#### Cloning of plasmid pRBPS-IVR7

In order to further suppress unintended expression of I-TevI which might have hampered the usability of pRBPS-IVR1, the lactose operator sequences (lacOs-spacer-lacOs) were located in front of the promoterless I-TevI expression cassette. In the resulting set-up the binding of LacI, (the repressor of the lactose operon) which is provided by the host strain, should have repressed possible background expression.

Therefore, the lacOs-spacer-lacOs element was removed from plasmid pRBPS-IVR1 and vector religation after removal of this fragment resulted in plasmid **pRBPS-IVR2**. Exactly the same fragment was reintegrated in plasmid pRBPS-IVR2 in front of the promoterless I-TevI expression cassette resulting vector **pRBPS-IVR3**. Therefore, vector pRBPS-IVR3 contains two symmetric lactose operator sites separated by a 147 bp spacer in front of the ITevI expression cassette. Although *lacI^q^* host strains were used which provided the repressor protein LacI, the insertion of a target sequence (e.g. an expression cassette) into the multiple cloning site of this construct was not possible.

To verify if the strong araBAD promoter reading counter clockwise towards the multiple cloning site hampered the integration of a target sequence, a vector was generated with the original orientation (referred to pRBPS7) of the arabinose P/O system including the ParA expression cassette. This was achieved by introducing the I-TevI restriction site into plasmid pRBPS7 in the first step and by introducing the I-TevI expression cassette including the minicircle part of pRBPS-IVR3 in the second step. The resulting vector of the first step, **pRBPS7_ITev-rs**, contains I-TevI restriction site and the parA expression cassette in the original orientation.

Plasmid pRBPS-IVR3 was used as the source of the I-TevI expression cassette and the minicircle core sequence (sequence between the resolution sites) and pRBPS7_ITev-res as the source for the arabinose P/O system including the ParA expression cassette and the I-TevI restriction sites. Fragments containing the described elements from both plasmids were generated. The extracted fragments derived from pRBPS-IVR3 and pRBPS7_ITev_rs were ligated resulting in vector **pRBPS-IVR4** containing the I-TevI expression cassette from pRBPS-IVR3 and the ParA expression cassette and the *araC* gene in the original orientation.

A further I-TevI restriction site was integrated in plasmid pRBPS-IVR4. A dsDNA fragment containing the 39 bp-Tev I site was generated by oligo annealing and integrated in vector pIVR4 resulting in plasmid **pRBS-IVR5** containing a second I-TevI restriction site.

The constitutive promoter of the *lacI* gene was inserted in plasmid pIVR5. This promoter was integrated in a way that it reads counter clockwise towards the I-TevI expression cassette and that the minicircle sequence of this parental plasmid is located between the promoter and the I-TevI expression cassette. Upon removal of the minicircle sequence by recombination the *lacI* promoter is situated close to the I-TevI expression cassette which can then be transcribed without hindrance. An appropriate sequence containing the lac promoter was designed for the insertion in plasmid pRBPS-IVR5 and ordered from Eurofins (Germany). Insertion of this sequence resulted in vector **pRBPS-IVR6** containing constitutive wt *lacI* promoter reading counter clockwise towards the I-TevI expression cassette until transcription is terminated by transcription terminator rrnbT1.

The wt lacI promoter was replaced by the lacI^q^ promoter. By exchange of a single nucleotide in the −35 bp region the promoter strength was enhanced by a factor ten according to the literature (Calos 1978). This point mutation was introduced in the wt lacI promoter sequence of plasmid pRBPS-IVR6 by site-directed mutagenesis. The nucleotide exchange was verified by sequence analysis. The resulting vector **pRBPS-IVR7** contains the lacI^q^ promoter reading counter clockwise towards the I-TevI expression cassette until transcription is terminated by transcription terminator rrnbT1.

### Insertion of reporter/target sequences into *in vivo* restriction vectors

#### Construction of vector pCMV-Luc

Vector pCMV-Luc was constructed by cloning the CMV promoter fragment of plasmid pcDNA3 (Invitrogen) into plasmid pGL2-Basic (Promega) using restriction sites *Mlu*I and *Hind*III. The resulting construct carries the wildtype luciferase gene (from *Photinus pyralis)* under the control of the immediate early promoter of the cytomegalovirus and the small t-antigen intron and the early poly(A) signal derived from simian-virus-40 at the 3’site.

#### Construction of plasmid pRBPS-IVR7_LucCMV

The Luciferase expression cassette of vector pCMV-Luc was subcloned into pRBPS plasmids as described by Mayrhofer (Mayrhofer et al. 2008). To construct plasmid pRBPS-IVR7_LucCMV, a wildtype firefly (Photinus pyralis) luciferase gene flanked by the immediate early CMV promoter at the 5’site and the SV40 small t-antigen intron & early poly(A) signal at the 3’ site was generated by digesting plasmid pRBPS7.CMV-luc with restriction enzymes *Spe*I and *Pst*I. The resulting fragment was cloned into the corresponding sites in plasmid pRBPS-IVR7 resulting in vector pRBPS-IVR7_LucCMV.

#### Construction of plasmidpRBPS-IVR7_BMP2

A PCR fragment comprising a BMP2 expression cassette containing promoter pEF1a and the polyA sequence derived from plasmid pEF1a-BMP2-advanced was generated using the primers EF1-B2s (5-ATTAGAGCTCATCTCGCTCCGGTGCCCGTCAGTG-3) containing a SacI site, EF1-B2as (5-TATTGGCCGGCCACGCCTTAAGATACATTGATGAGTTTG −3’) containing a FseI site and vector pEF1a-BMP2-advanced as template. Plasmid pEF1a-BMP2-advanced was described by Hacobian et al (Hacobian et al. 2016). The 50 μl PCR reaction containing 2.5 μl (10mM) primer DNA, 1μl (10 mM) dNTPs, 20 ng template DNA, 1.5μl DMSO, 0.5 μl Phusion polymerase in HF buffer was subjected to the following conditions: 30 sec 98°C pre-denaturation, 40 cycles: 30 sec 98°C, 20 sec 61°C, 90 sec 72°C. The PCR product was inserted blunt into plasmid pCR-Blunt. A SacI fragment was extracted from the resulting vector which was cloned into the corresponding single restriction site of pRBPS-IVR7 resulting in vector pRBPS_IVR-BMP2.

### Lab scale minicircle-DNA production

For lab scale production either 100 ml or 1000 ml cultures of parental plasmid pRBPS-IVR7_LucCMV carrying the Luciferase expression cassette under the control of the CMV immediate early promoter were cultivated. LB -medium supplemented with kanamycin were inoculated with 1/100 volume of pre-culture, i.e. 1 ml or 10 ml of an overnight culture of an *E. coli* strain transformed with plasmid pRBPS-IVR7_LucCMV grown at 28°C in LB-medium supplemented with kanamycin and 1% glucose. The cultures were cultivated at 37°C and the expression of the ParA resolvase was induced with 0.5 % L-arabinose. For optimal results in shaking flask cultures induction was accomplished at an OD600 range of 1.5 to 2. The resulting minicircle-DNA was purified using standard kits for pDNA preparation.

### Industrial pilot scale production of BMP2 minicircle-DNA using parental plasmid pRBPS-IVR7_BMP2

On day 1 two independent pre-cultures were prepared by inoculating 0.2 μl and 2 μL of *E. coli* BL21 stock harbouring plasmid pRBPS-IVR7_BMP2 in 2L shake flasks each containing 500 mL of LB medium supplemented with 1 % D-Glucose and 50 mg/L kanamycin sulphate. The pre-cultures were grown overnight for about 16 hours at 28°C in a NBS INNOVA 42R incubator at a shake speed of 270 rpm. The next day (day 2, morning) seeds from the most appropriate pre-culture were inoculated to reach a starting OD600 = 0.006 into 4 Sartorius biorectors (5L working volume) each containing 4.5L of PPM medium supplemented with 50 mg/L kanamycin sulphate. The fermentors were operated with strictly controlled parameters including pH, temperature and dissolved oxygen (regulated by cascades on airflow, agitation and pressure). The cultures were grown for 6 hours at 28°C before the medium fed-batch started. The temperature was then raised (day 3) to 37°C according to a 30 minutes linear temperature shift ramp after 28, 25, 22 or 19 hours growth at 28°C respectively and biomass production phases at 37°C lasted for 3, 6, 9 and 12 hours respectively. Following the temperature shift each culture was supplemented with 10g/L arabinose to induce the expression of the ParA resolvase (ParA) leading to the production of the BMP2 minicircle. Induction was performed for three hours at 37°C. Cells were harvested at 4°C and 7000 RPM for 25 min using a Beckman J-26 XP centrifuge and a Avanti JLA 8.1000 rotor. The pellets were then stored in bags at −20°C. To analyse the fermentation process and outcome, plasmid DNA was purified at various time points using standard miniprep kits (QIAprep Spin Miniprep kit, Qiagen, Germany).

### Large scale production of luciferase minicircle-DNA using parental plasmid pRBPS-IVR7_ LucCMV

Parental plasmid pRBPS-IVR7_LucCMV was tested for large scale production using the best culture condition identified previously for the pRBPS-IVR7_BMP2 plasmid (see results). Briefly, the same pre-culture and biomass production at 28°C as for the pRBPS-IVR7_BMP2 candidate were used and followed by a 3 hours biomass production and a 3 hours induction phases at 37°C. This production was done in duplicate.

### Cell lysis and Clarification

Bacterial lysis was performed following the procedure described in CIMmultus™ HIP2 Plasmid Process Pack™ (Publication#: IM-CIMacpDNA-1203; BIA Separations, Ajdovščina, Slovenia, 2012). The culture of *E. coli* containing N-RBPS-LUCIF-F01 prepared by Eurogentec was thawed and re-suspended in 50 mM TRIS – HCl buffer pH 8.0 containing 10 mM EDTA. When the suspension was homogenous it was treated with 0.2 M NaOH, 1% SDS lysis buffer for 5 min, followed by addition of neutralization buffer, 3.0 M CH3COOK pH 5.0. Then 5 M CaCl2 was slowly added to the final concentration of 0.75 M. Such solution was incubated for 15 min while gently mixing it. The mixture was coarse filtrated through cotton gauze and strainer and then fine filtrated with Sartopore® 2 0.45um (Sartorius AG, Goettingen, Germany) filter into clean vessel

### Chromatographic Purification

#### Ion Exchange Chromatography

For the first chromatographic step, the capture step, a weak anion exchanger CIMmultus^TM^ DEAE-800 (BIA Separations, Ajdovščina, Slovenia) was used. The conductivity of fine filtrated material was adjusted to 35 mS/cm with addition of sufficient amount of deionized water filtrated with PES 0.22μm filter (TPP, Trasadingen, Switzerland). Material was loaded with flowrate of 800 ml/min and step elutions were performed with 400 ml/min. Buffers used for capture step chromatography were 50 mM TRIS – HCl, 10 mM EDTA, pH 7.2 as equilibration buffer and 50 mM TRIS - HCl, 10 mM EDTA, pH 7.2 with 0.6 M or 1 M NaCl as elution buffers. Absorbance at 260 nm was monitored during chromatography run. At the end of each run the column was washed with 1 M NaOH and subsequently with 2 M NaCl to clean and regenerate the resin.

#### Hydrophobic Interaction Chromatography

For the polishing step CIMac™ C4 HLD and CIMmultus™ C4 HLD-8 columns (BIA Separations, Ajdovščina, Slovenia) were used. After finding the right conditions for optimal purity by using linear gradient a sample displacement chromatography mode was implemented. This technique results in better purity of the final product by displacing other DNA forms from the column.

To perform the polishing step the main elution fraction from the capture step was applied after dilution with 50 mM TRIS – HCl, 10 mM EDTA, 4 M (NH4)_2_SO_4_, pH 7.2 to the final loading concentration of 1.7 M ammonium sulphate. subsequently the material was loaded until the second break through, followed by washing the column thoroughly with 50 mM TRIS – HCl, 10 mM EDTA, 1.7 M (NH_4_)_2_SO_4_ pH 7.2 buffer. Elution steps were performed with 50 mM TRIS – HCl, 10 mM EDTA, pH 7.2 and decreasing (NH_4_)_2_SO_4_ concentrations to 1.5 M, 0.8 M and finally to 0 M. All chromatographic fractions were analyzed either by agarose gel electrophoresis or HPLC analytics.

### Analytics of fractions

Fractions from preparative runs were analyzed with HPLC analytics using PATfix™ (BIA Separations, Ajdovščina, Slovenia) with a 50 mm biocompatible optical cell or a Äkta Purifier 10 system (GE Healthcare Life Sciences, Boston, United Sates) with 10 mm optical cell and a CIMac™ pDNA-0.3 Analytical Column (BIA Separations, Ajdovščina, Slovenia). Buffers used were 200 mM TRIS – HCl, 10 mM EDTA, pH 8 with or without 1 M NaCl. Separation of different DNA forms was achieved by applying a linear gradient from 0.5 M to 0.8 M NaCl or from 0.6 M to 0.7 M NaCl in 10 min.

For analytics by gel electrophoresis 1% agarose gels were used in a horizontal electrophoresis unit. Mass Ruler DNA ladder-mix of 10 kbp and concentration of 103 ng/μl was applied (Thermo Fisher Scientific, Massachusetts, USA). Analysis of the gel was performed at SyngeneGiBox, using Gensys software.

### In vitro assays

#### Cell Culture

For in vitro analysis, Chinese hamster ovary cells (CHOs, ATCC® CCL-61TM, USA) were used (DSMZ, Braunschweig, Germany, #ACC565). Cells were grown in Dulbecco’s modified Eagle’s medium high glucose (DMEM high glucose; Sigma-Aldrich, USA) supplemented with 10 % FCS (Lonza, Basel, Switzerland) and 1 % L-glutamine (Sigma-Aldrich, USA) at 37 °C and 5 % CO2. 0.4 × 10^5^ cells/well were seeded in a 24-well plate and transfected at approximately 80 % confluence using equimolar as well as same amount (1 μg) of DNA (either minicircle-DNA or conventional plasmid DNA) and jetPEI® DNA transfection reagent (VWR, Pennsylvania, USA; 1:2 v/v) as recommended by the manufacturer. For in vitro assay testing minicircle-DNA produced according to the lab scale protocol was used.

#### Luciferase Assay

The enzymatic activity was measured 48 h, 1 week, 2 weeks and 3 weeks after transfection in CHO cells. Cells were washed with 100 μl PBS and lysed with 100 μl cell lysis solution (Promega, Madison, USA). 10 μl of samples were incubated with 100 μl luciferase assay reagent (Promega, Madison, USA) for quantification in a 96-well plate.

### QC analysis

The minicircle produced in this study were subjected to several quality control assays that meet ICH compliance (ICH), namely topology analysis by HPLC, endotoxins content by LAL kinetic assay, residual host cell DNA by qPCR, residual host cell RNA by HPLC and host cell protein content by ELISA.

#### Quantification of endotoxins

The endotoxin content was measured by the chromogenic limulus amoebocyte lysate method (KQCL) using theKinetic-QCL™ Chromogenic LAL Assay kit from Lonza.

#### Quantification of host cell RNA

Quantification of residual RNA was performed by HPLC analysis. Briefly, a TSKgel® OligoDNA RP column (Tosoh Bioscience, ref. 13352) filled with spherical 5 μm particle with 250 Å pore size, C18 derived silica-based matrix specifically designed for the purification of oligonucleotides, RNA and DNA fragments (up to 500mer) was used. The column, thermostated at 40 °C, was utilized to separate plasmid DNA and RNA using a linear acetonitrile gradient (0 to 40%) over 24.05 minutes. RNA standards in the range of 4 to 44 μg/ml were prepared from *E. coli* NTC4862 cells using the High Pure RNA Isolation kit from Roche Sample material and standard solutions were digested with RNase solution (at 1 U for 100 μg of RNA) for 10 minutes at room temperature. Samples were analysed by HPLC using an Agilent 1260 Infinity II HPLC system. The proportional linear relation between the concentrations of RNA standards and their peak areas obtained in the chromatograms was used to calculate the residual RNA content in the sample material.

#### Quantification of residual host cell DNA

The residual host cell DNA content in the bulk material was measured by a quantitative polymerase chain reaction assay as described by Lee et al (Lee et al. 2010).

#### Quantification of Host Cell Protein

Quantification of residual host cell protein (HCP) was performed using E.coli HCP ELISA kit from Cygnus technologies LLC (Southport, USA).

#### Topological analysis

Isoforms of minicircle-DNA were identified by CGE and HPLC. Briefly, HPLC analysis were done using Tosoh Bioscience TSKgel DNA-NPR weak anion exchange column (Tososh Biosciences, ref# 18249) coupled with a DNA-NPR guard column (Tososh Biosciences, ref# 18253). Buffers used were 25 mM sodium borate pH 9.00 with or without 1M NaCl. Separation of different isoforms was achieved by applying a linear salt gradient form 0.55 M to 0.75 M over 2 column volumes.

As for the GCE analysis samples were analysed using a Beckman PA800 Plus CE instrument equipped with an integrated solid state 488 nm laser light source (Beckman, Paolo Alto, CA, USA) for laser-induced fluorescence detection. The wavelength of excitation was 491 nm and the emission fluorescence wavelength was set to 509 nm. Separation was performed in 100-μm-i.d. coated capillaries (Sciex, Toronto, Canada) with a total length of 40 cm. Electrophoresis was performed with negative source polarity at 250V/cm at 30°C. Plasmid samples were applied by pressure injection with an overpressure of 0.1 psi for 1s, followed by an injection of run buffer for 1 s. Prior to injection of a sample, the capillary was equilibrated with a 7× dilution gel solution in TBE 1× buffer (Sigma-Aldrich, USA) containing the intercalating dye YOYO-1 (1.1 v/v) (Thermo Fisher Scientific, Massachusetts, USA). The data were analyzed using 32 Karat Software (Beckman). For relative quantification of the different plasmid DNA isoforms, the integrated peak areas were calculated according to the corrected area percent method.

Standards for the topological analysis were prepared as follows: Open circular (OC) isoforms were generated from purified samples of fermentation run LUCIF-F01 and BMP2-F09 using nicking endonuclease Nt.BsmAI while linear (L) forms were prepared using endonuclease XhoI and PsTI for luciferase and BMP2 minicircles respectively according to the manufacturer’s instructions. The OC and L isoforms were then purified using a Maxiprep kit and the quality was confirmed by agarose gel electrophoresis, DNA-NPR HPLC and CGE analysis as described above.

## RESULTS

### Parental Plasmid pRBPS-IVR

The goal of this work was the development a plasmid that enables the coordinated, consecutive performance of *in vivo* processes involved in minicircle production and to prove the scalability of this system for industrial large scale manufacturing. To achieve this, the elements on the parental plasmid were arranged in a way that expression of the endonuclease needed for degradation was activated through the removal of the minicircle region by site specific recombination. The detailed set-up of this novel minicircle production vector is shown in Figure 2. This plasmid carries the ParA resolvase (ParA) under the control of the arabinose promoter/operator system (Pbad, ParaC, araC), a promoterless expression cassette of the ITev endonuclease (ITev, SD), ITev restriction sites (ITev-RS), symmetric versions of lactose operator sites (lacOs) separated by spacer and located in the miniplasmid region (non-therapeutic region), the resolution sites (res) for site-specific recombination, the terminators T1 and T2 located within the minicircle region (i.e. between the res sites), the constitutive promoter lacIq orientated counterclockwise towards the ITev expression cassette, as well as the aminoglycoside 3’-phosphotranserase sequence conferring resistance to kanamycin (Kan) and the MB1 origin of replication (ORI).

**Fig. 2.**
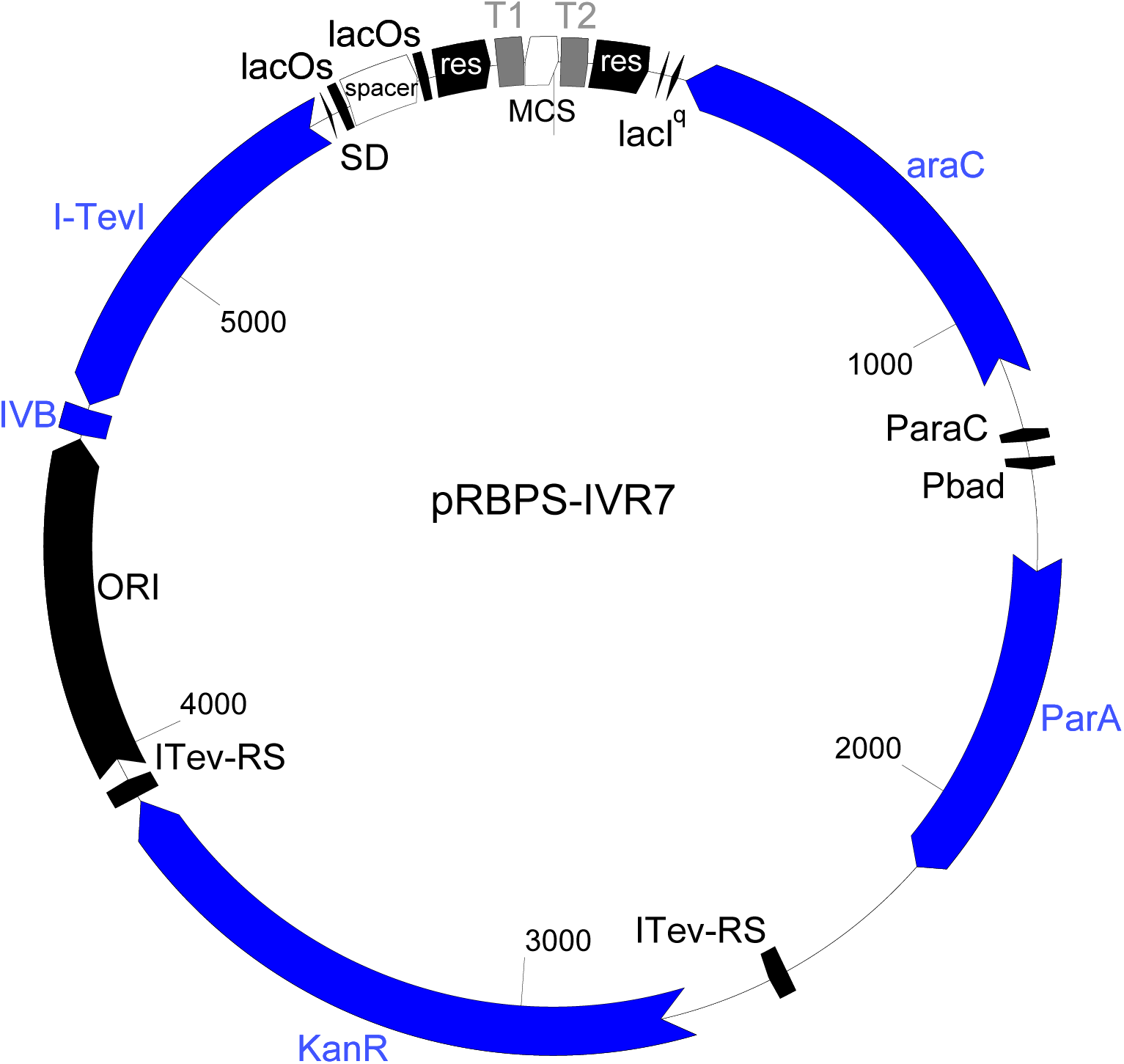
Map of plasmid pRBPS_IVR7: T1, transcription terminator 1; MCS, multiple cloning site, T2, transcription terminator 2; res, resolution site of the ParA resolvase system; lacIq, −35 and −10 region of an up-mutation of the constitutive promoter of the lacI gene of the lactose operon; araC, gene enconding the repressor of the arabinose operon; ParaC, promoter of the araC gene; Pbad, promoter of the araBAD genes of the arabinose operon; ParA, parA resolvase gene; ITev-RS, ITev restriction site; Kan, aminoglycoside 3’-phosphotransferase expression cassette conferring resistance to kanamycin; ORI, MB1 origin of replication; ivb, *in vivo* biotinylation sequence; ITev, intron enconded endonuclease of bacteriophage T4; SD, Shine Dalgarno sequence for ITev translation; lacOs, modified lactose operator site; spacer, spacer sequence between two direct repeats of the modified lactose operator sites lacOs

In this set-up, the expression of the endonuclease is initiated by the recombination step for minicircle-DNA production. Basically, the counter clockwise orientated lacIq promoter is separated from the ITev expression cassette by the minicircle region of the parental plasmid. As shown in Figure 2, terminators T1 and T2 are integrated in this region. Expression of the ParA resolvase, which is triggered by the addition of L-arabinose, mediates the recombination process. Consequently, all sequences, including the terminators T1 and T2, located between the resolution sites are excised resulting in the desired minicircle-DNA. Prior to the recombination event, transcription from the lacIq promoter is terminated by terminators T1 or T2 at latest. After removal of these sequences by recombination, the lacIq promoter drives the transcription towards the ITev expression cassette without hindrance, thereby producing ITev mRNA. Following expression, the ITev endonuclease acts at the corresponding restriction sites (ITev-RS) thereby linearizing miniplasmid-DNA, which is subsequently further degraded by host exonucleases. This system set-up guarantees the coordinated execution of recombination and restriction, as expression of the endonuclease depends on the successful performance of the recombination step.

### Fermentation/large scale production

Two different construct were investigated to test the scalability of pRBPS-IVR system, namely plasmid pRBPS-IVR7_BMP2 carrying an expression cassette for bone morphogenetic protein 2 and plasmid pRBPS-IVR7_LucCMV carrying an expression for the reporter gene firefly luciferase. Compared to lab scale production, prolonged periods of biomass production (20 hours and more) are necessary in large scale approaches before the desired optical density of the culture is reached to induce the recombination/degradation process. In order to optimize stability of parental plasmids during this period, cultivation temperature was decreased to 28°C. At this temperature the plasmid copy number established by the pMB1 origin of replication is lower than at 37°C and additionally, the arabinose promoter system is more tightly repressed. With regard to product yield the high copy number situation at 37°C needs to be re-established before the recombination/degradation process is induced. To determine the time necessary at 37°C to return to a high copy number phenotype, four different periods at this temperature were tested with E.coli BL21 (pRBPS-IVR7_BMP2) in fed-batch fermentations (F1 to F4) at 4.5 l scale (+ feed). A detailed fermentation scheme is depicted in Figure S2 of the supplementary material.

Fermentation runs with *E. coli* BL21 harbouring plasmid pRBPS-IVR7_BMP2 were performed as described in materials and methods under “Industrial pilot scale production of BMP2 minicircle-DNA using parental plasmid pRBPS-IVR7_BMP2”. Recombination and subsequent degradation were induced by addition of 1 % L-arabinose at an optical density of approximately 60 and the culture reached an OD_600_ of about 80 in case of pRBPS-IVR7_BMP2. Comparable results were achieved with plasmid pRBPS-IVR7_LucCMV under the same conditions. However, as stated in Table 1 the final yield was higher in case of the BMP2 construct were about 108 μg/ml have been achieved in this initial trials.

**Table 1.**
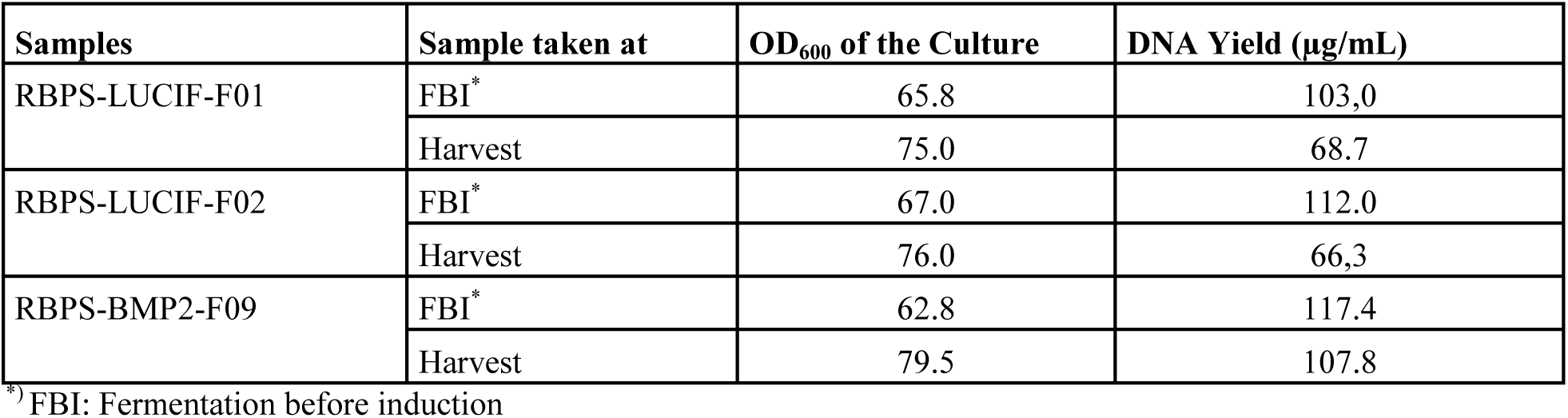
Summary fermentation yield.

After fermentation, an aliquot of each run was taken and subjected to plasmid DNA isolation and further analysis by restriction digestion and AGE (data not shown). These results indicated that the parental plasmid was correctly produced and stable at 28°C. Prolonged cultivation at 37°C however, led to gradually degradation of the parental plasmid indicating that this phase, which is needed to achieve high plasmid copy number, should be kept as short as possible but as long as needed to establish the high copy number phenotype. As stated in materials and methods time frames of 12, 9, 6 and 3 hours at 37°C were tested. The results achieved showed that 3 hours at 37°C were sufficient to increase the copy number and did not affect the integrity of the parental plasmid.

The results achieved with pRBPS-IVR7_BMP2 were applied to a further parental plasmid to see whether they were specific for the tested construct or transferable to other minicircle producing RBPS parental plasmids. The next parental plasmid tested in fermentation runs was pRBPS-IVR7_LucCMV. Applying the same conditions as with the BMP2 construct resulted in the same fermentation profile and stable propagation of the parental plasmid with no visible degradation before induction. The result of these fermentation runs can be seen in Figure 3 showing an AGE analysis of the products.

**Fig. 3.**
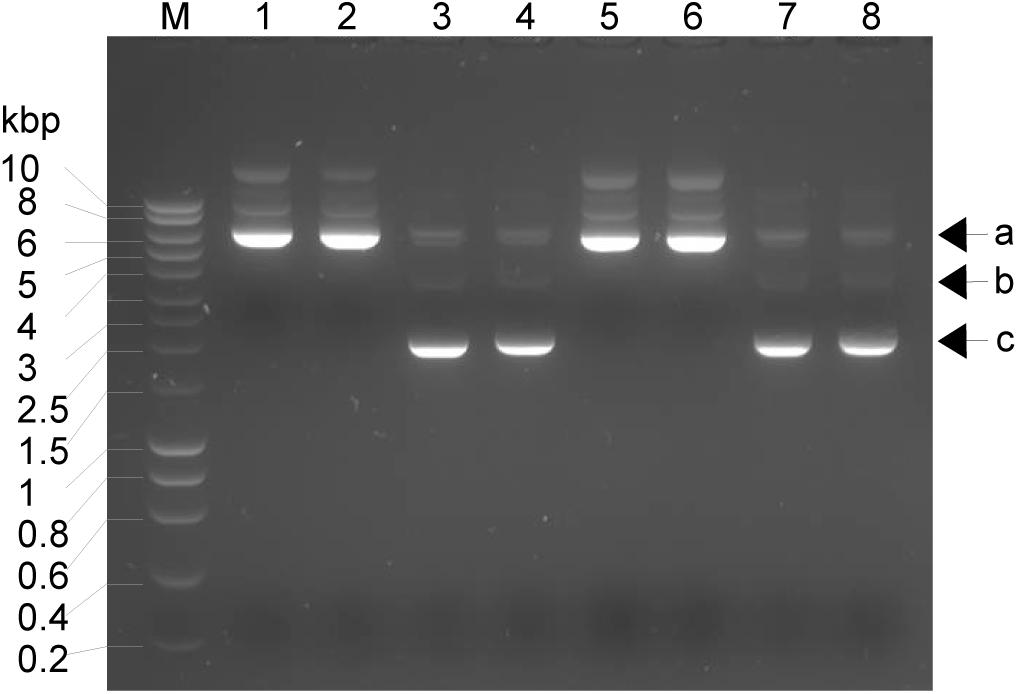
shows AGE analysis of fermentation runs of E.coli BL21 (pRBPS-IVR_LucCmv) which were performed in duplicate (F01, F02). Samples were taken at various time points during fermentation runs F01 and F02, pDNA was isolated and subsequently subjected to AGE. lane M, marker; lane 1, F01 2 hours 05 min. after shift to 37°C; lane 2, F01 at the time point of induction; lane 3, F01 1 hour 30 min. after induction; lane 4, F01 at the time point of cell harvest; lane 5, F02 2 hours 05 min. after shift to 37°C; lane 6, F02 at the time point of induction; lane 7, F02 1 hour 30 min. after induction; lane 8, F02 at the time point of cell harvest; a, (residual) parental plasmid supercoiled form and minicircle-DNA; b, minicircle-DNA linear form; c, minicircle-DNA supercoiled form

According to Figure 3, in terms of minicircle yield and residual parental plasmid it seems that 1.5 hours of induction gives a result very comparable to that achieved after 3 hours of induction. Only minimal amounts of residual parental plasmid can be observed after both induction periods.

### Purification

The downstream processes applied to purify minicircle-DNA produced with the RBPS *in vivo* restriction technology are basically very similar to standard processes used for conventional plasmids. Despite the recombination efficiencies achieved in this first fermentation trials a minor fractions of residual parental plasmid was still contaminating the product. According to the goal of this study a scalable process was developed to remove this contamination. To achieve this, a combination of conventional chromatographic techniques namely ion exchange and hydrophobic interaction chromatography was applied to the samples produced in the described fermentation runs. The data for the luciferase minicircle-DNA purification process are presented below.

After cell lysis and clarification a sample capture step was performed using a weak anion exchanger. The sample was loaded onto a CIMmultus™ 800 column and eluted as described in materials and methods.

The loaded sample as well as further relevant fractions of the anion exchange chromatography run shown in Figure 4a have been analysed by HPLC and AGE. Figure 4b shows an overlaid chromatogram from HPLC analysis of the loaded sample, elution fraction 3 (E3) and the tail of elution fraction 3 (E3t). Figure 4c shows the analysis of the loaded sample, the flowthrough and the elution fractions 1 to 3 by AGE. Due to the high salt concentration in the elution fractions the DNA shows different migration behaviour in AGE. The minicircle consists of 3.735 bp and its supercoiled band should therefore be expected around the 2.5 kbp marker band whereas in Figure 4c the band appears between 5 and 6 kbp. Summing up Figure 4 shows that the loaded sample contained the minicircle-DNA but also RNA and minor amounts of residual parental plasmid. Most of the RNA eluted in the flowthrough and in elution fractions 1 and 2, whereas no minicircle-DNA can be observed in these fractions. Elution fraction 3 and 3t contained the minicircle-DNA and residual parental plasmid. The overlaid chromatograms of fractions L, E3 and E3t shows two peaks whereas the smaller, earlier one was attributed to the residual parental plasmid. Interestingly, in the elution fraction E3 of the capture step the ratio between minicircle-DNA and residual parental plasmid obviously shifted in favour for the minicircle-DNA.

**Fig. 4.**
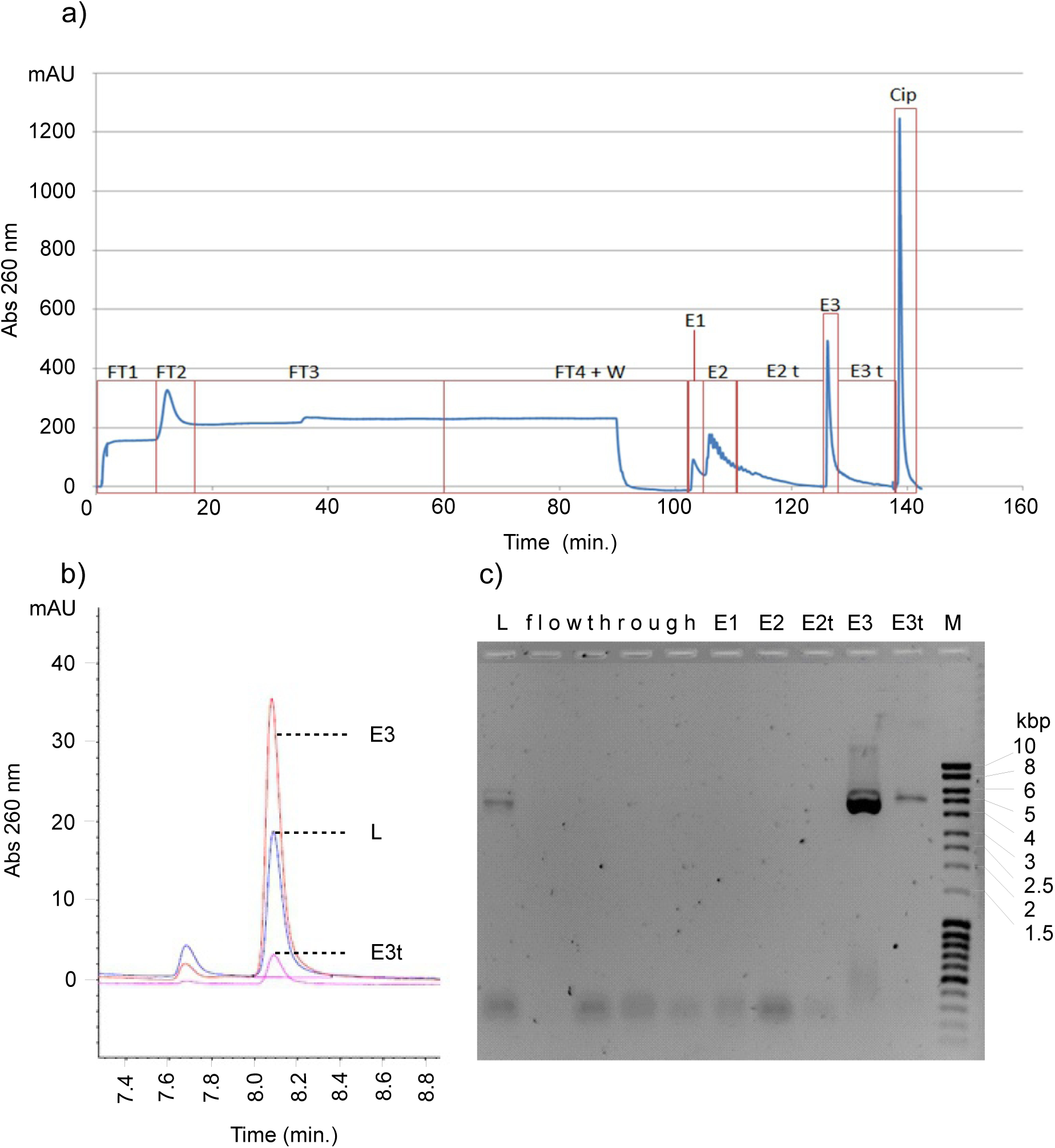
shows the results of the chromatographic capture step. **Fig. 4a)** The chromatogram of the ion exchange run using CIMmultus™ DEAE-800 is shown. Flowrate for loading was 800 ml/min and for elutions 400 ml/min. Elution steps were performed with TRIS – HCl buffer with 0.6 M and 1 M NaCl. The flowthrough fractions are indicated as FT1 to FT4. The elution fractions are marked with E1 to E3 whereas a tail fraction was collected in case of elution fraction 2 and 3 designated as E2t and E3t. Cip is the peak resulting from the cleaning in place procedure. **Fig. 4b)** The chromatogram of the HPLC analysis of fractions E3, E3t and the loaded sample (L). **Fig. 4c)** AGE analysis of the indicated fractions from the ion exchange chromatography step: L, loaded sample; flowthrough, fractions FT1, FT2, FT3 and FT4 shown in a); E1, elution fraction 1; E2, elution fraction 2; E2t, tail of elution fraction 2; E3, elution fraction 3; E3t, tail of elution fraction 3; M, marker

After successful capturing of the minicircle-DNA by anion exchange chromatography, a strategy to remove residual parental plasmid was developed. This strategy was based on hydrophobic interaction chromatography (HIC). First analytical trials to explore conditions appropriate to separate the parental plasmid from the minicircle were performed on CIMac™ C4 HLD analytical column (BIA Separations, Ajdovščina, Slovenia) with aliquots of elution fraction 3 from the anion exchange capturing step shown in Figure 4. HIC was performed using the buffer described in materials and methods by applying an ammonium sulphate gradient from 3 M to 0 M. Based on these results the conditions for sample displacement were calculated and the polishing step in the sample displacement mode was implemented, again using a CIMac™ C4 HLD analytical column. Step elution was performed as described above.

Figure 5 shows the HIC run in displacement mode using a CIMacTM C4 HLD analytical column. This figure shows the first breakthrough at approximately 4 ml and the second at approximately 7 ml. Further analysis by HPLC and AGE of fractions shown in Figure 5 is presented in the supplementary material (Fig. S3).

**Fig. 5.**
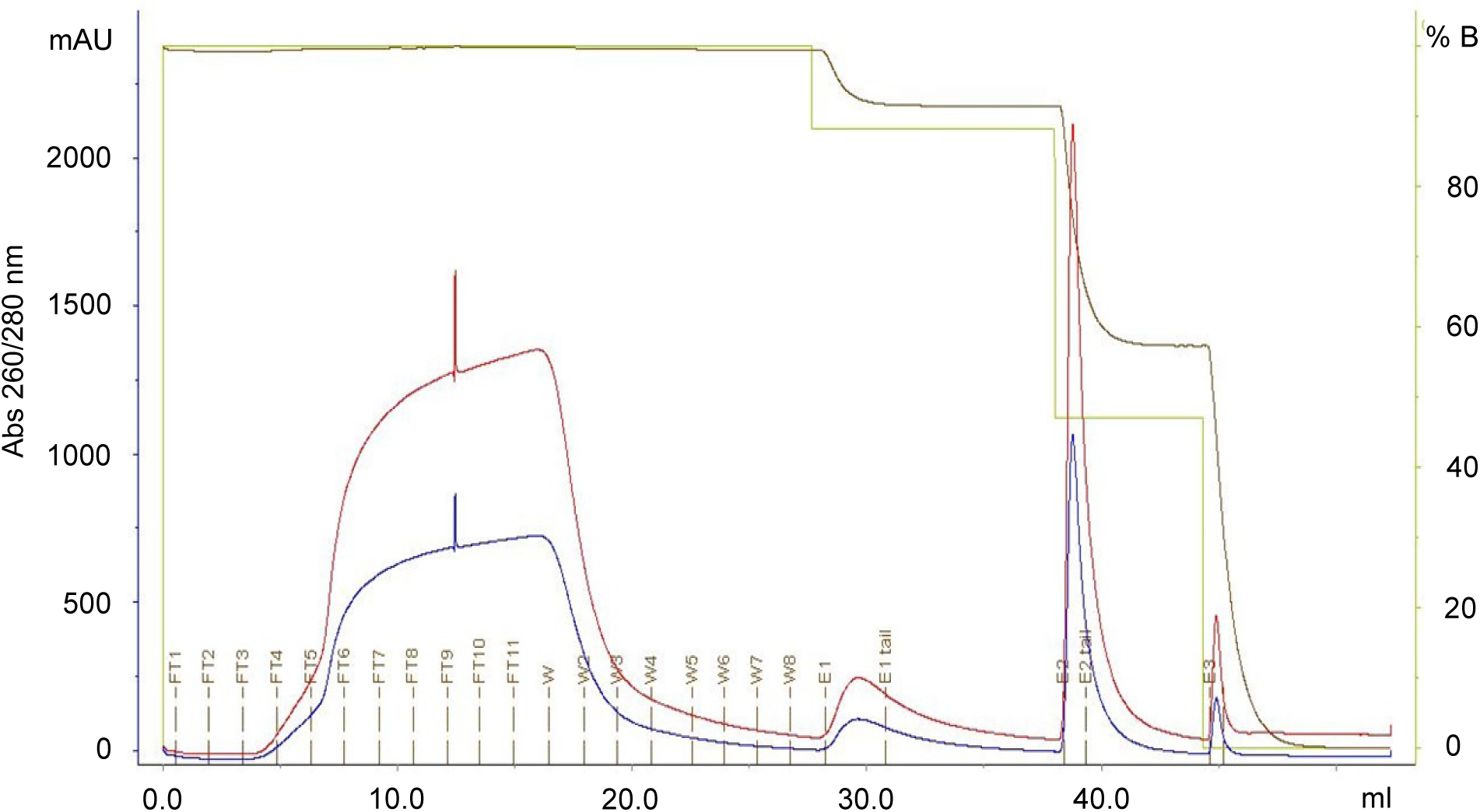
shows the Chromatogram of the analytical sample displacement chromatography implemented on CIMac^TM^ C4 HLD column. The flowrate for loading, washing and elution was 1 ml/min. The signals monitored were absorbance at 260 nm (red), absorbance at 280 nm (blue), conductivity (brown) and concentration of pump B - 50 mM TRIS – HCl, 10 mM EDTA, 1.7 M (NH_4_)_2_SO_4_ pH 7.2 buffer (green). Material was loaded in 50 mM TRIS – HCl, 10 mM EDTA, pH 7.2 buffer with final concentration of 1.7 M (NH4)2SO4, followed by washing the column in the same buffer. Step elution was performed with 50 mM TRIS – HCl, 10 mM EDTA, pH 7.2 buffer with final concentrations of ammonium sulphate of 1.5 M (E1), 0.8 M (E2) and 0 M (E3). FT1 - FT11, flowthrough fractions; w1 - w8, washing fractions; E1 - E3, elution fractions

The sample displacement strategy is based on the fact that under certain conditions the minicircle-DNA binds the matrix stronger than the parental plasmid. Under such conditions the minicircle-DNA replaces already bound parental plasmid from the chromatographic matrix if sample is loaded to the column beyond its capacity.

Scale up from analytical to preparative scale was performed using the same sample from the capture run (fraction E3) and a CIMmultus™ C4 HLD-8 column. Running conditions were basically the same except of the flowrates, which were adjusted to 15 ml/min for loading and 10 ml/min for elution. Using this strategy, we finally managed to elute highly pure minicircle-DNA in fraction E2 as demonstrated by AGE and HPLC analysis (Figure 6a and 6b) whereas sample purity based on HPLC analytics was calculated to 99%. The AGE analysis of the product supports the result achieved by HPLC. Based on the CGE result shown in Figure 6c a supercoiled content of 92% was calculated (see QC results). Summing up, the results presented in this report strongly suggest that mc-DNA production based on the next generation of RBPS parental plasmids introduced herein is scalable and can be used for large scale GMP production.

**Fig. 6.**
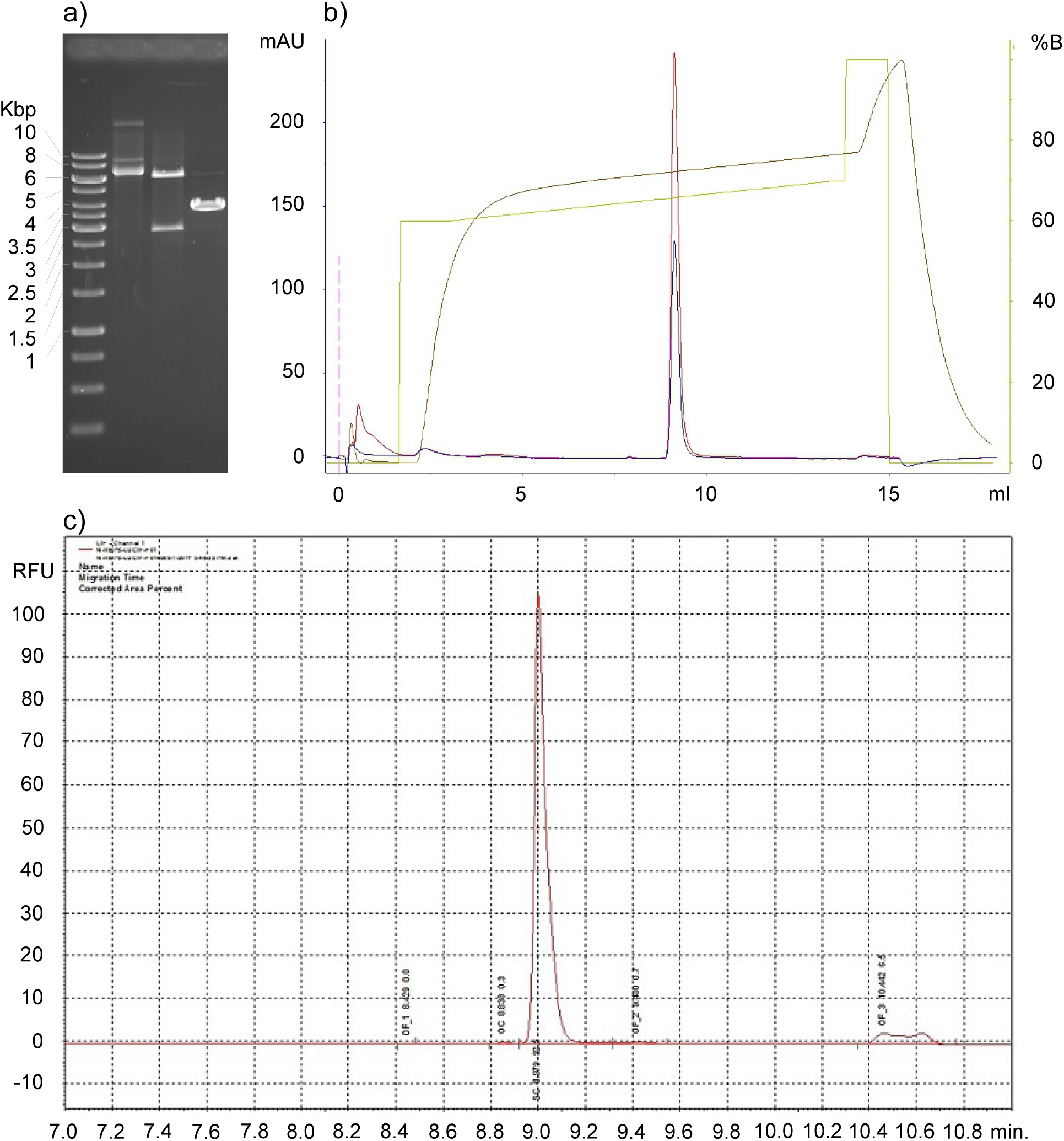
shows AGE and HPLC analysis of the purified luciferase minicircle-DNA. **Fig. 6a)** Shows restriction analysis and AGE of the parental plasmid pRBPS-IVR_LucCMV and the final product, the luciferase minicircle-DNA. Both samples were digested with Hindlll resulting in a 6.415 bp and a 2.887 bp fragment in case of parental plasmid pRBPS-IVR7_LucCMV and in linearization of the luciferase minicircle-DNA (3.735 bp). **lane 1**, parental plasmid pRBPS-IVR7_LucCMV undigested; **lane 2**, parental plasmid pRBPS-IVR7_LucCMV digested with Hindlll; **lane 3**, luciferase minicircle-DNA digested with Hindlll. **Fig. 6b)** Shows a Chromatogram of fraction E2 from the scale up run on CIM™ C4 HLD-8, analysed by HPLC. The signals monitored were absorbance at 260 nm (red), absorbance at 280 nm (blue), conductivity (brown) and concentration of pump B. The x-axis shows the elution volume in milliliter (ml), the y-axis shows the absorbance in milli absorbance units (mAU). The peak close to 10 ml indicates the supercoiled luciferase minicircle-DNA. **Fig. 6c)** Shows an electropherogram of fraction E2 from the scale up run on CIMTM C4 HLD-8. The x-axis shows the retention time in minutes (min.), the y-axis shows the relative fluorescence units (RFU). The peak at 9 minutes indicates the supercoiled luciferase minicircle-DNA making up 92.5% of the total DNA shown in this electropherogram

### In vitro assays Luc minicircle vs Luc reference plasmid

In the course of this study experiments were performed to check how minicircle-DNA produced with the RBPS *in vivo* restriction technology would perform in gene transfer experiments compared to conventional pDNA. For this purpose CHO cells were transfected with minicircle-DNA and conventional DNA both coding for the same expression cassette for the enzyme firefly luciferase using the jetPEI® transfection reagent. The enzyme activity was measured after 48 hours, 1 week, 2 weeks and 3 weeks using an assay kit for quantification in 96-well plates. The experiments were carried out to compare both plasmid species at an equimolar level corresponding to 1μg (256 pmol) of conventional plasmid DNA (Figure 7a) and at different levels of minicircle DNA (Figure 7b) using either 1 μg minicircle DNA or an amount corresponding to 1μg of the conventional plasmid (i.e. 0.6μg minicicircle DNA). Each experiment shown in Figure 7 was performed in duplicates and each value is an average of a triple measurement.

**Fig. 7.**
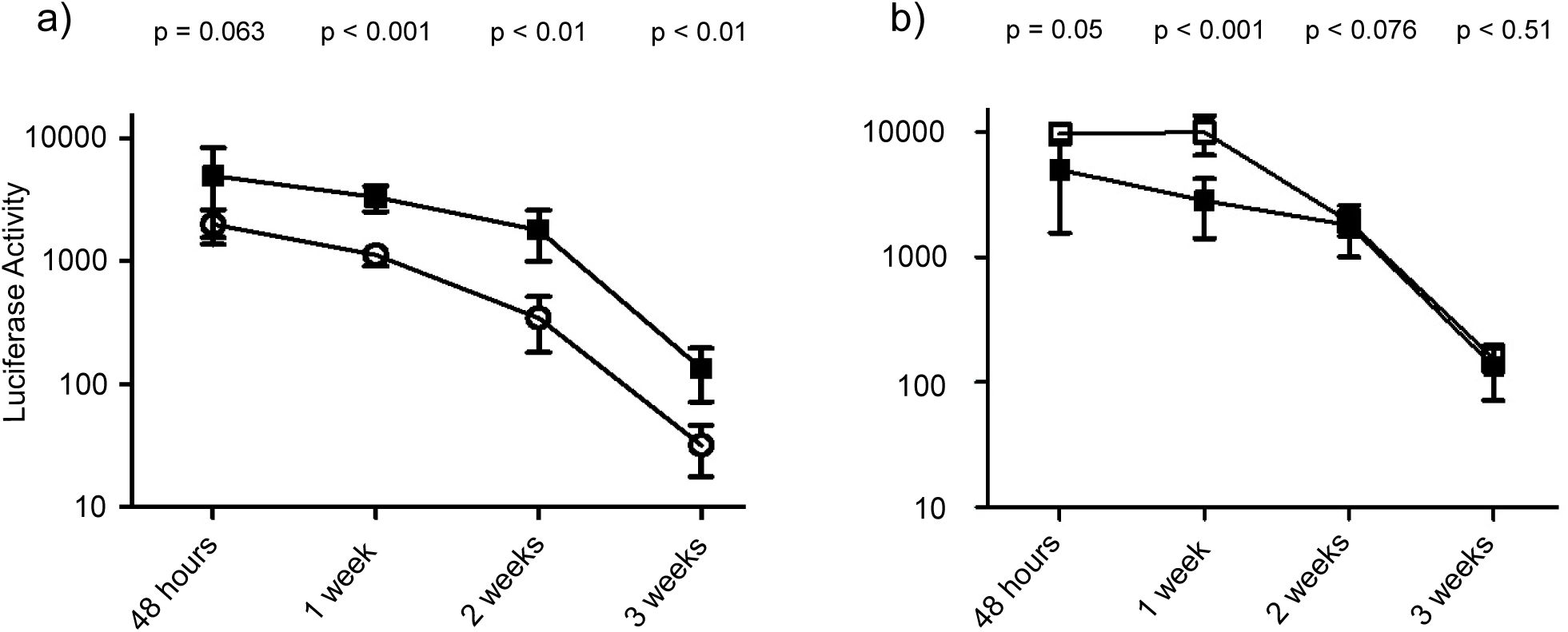
shows the results of a luciferase activity assays performed with either minicircle-DNA (MC-Luc) or with a conventional plasmid (pCMV-Luc) encoding the same luciferase expression cassette. **Fig. 7a** Minicircle and conventional plasmid DNA were used in equimolar amounts (256 pmol) corresponding to 1 μg of the conventional pDNA. CHO cells were transfected using the jetPEI® DNA transfection reagent. The enzymatic activity was measured after 48 hours, 1 week, 2 weeks and 3 weeks as indicated on the x-axis of the figure. 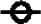: pCMV-Luc; 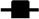 MC-Luc. **Fig. 7b** shows the result of a luciferase activity assay performed with different amounts of minicircle-DNA (MC-Luc). In this assay the effectivity of different amounts of minicircle-DNA was compared. The time series were either performed with 1 μg of minicircle-DNA, or with an amount (256 pmol, 0.6 μg) corresponding to 1 pg of the conventional plasmid pCMV-Luc carrying the same expression cassette. CHO cells were transfected using the jetPEI® DNA transfection reagent. The enzymatic activity was measured after 48 hours, 1 week, 2 weeks and 3 weeks as indicated on the x-axis of the figure. 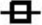 : MC-Luc lμg; 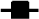 MC-Luc 0.6μg (256 pmol)

As demonstrated in Figure 7a the minicircle-DNA showed significant stronger luciferase activity at every time point measured. There is only a moderate decrease within the first two weeks and even after three weeks the activity is still in a range comparable to week 2 of the conventional plasmid. Using the same total amount as with the conventional plasmid DNA (i.e. 1 μg), the minicircle of course demonstrated even higher enzyme activity at the first two time points measured. Interestingly, after two weeks the enzyme activity induced by 1 μg minicircle-DNA was in the same range as the activity induced by 0.6 μg (the amount equimolar with 1 μg conventional pDNA). This indicates that for long term activity of a DNA agent administered as minicircle-DNA the lower dose (equimolar to the conventional pDNA) is sufficient and levels out to the same effect as the higher dose of minicircle DNA. This indicates clearly that there is no advantage associated with the application of the higher dose.

### QC Results

The produced minicircle-DNA was analysed to check for quality requirement as demanded by health care authorities. The results are summarized in Table 2.

**Table 2.** Summary quality control results.

| Impurity Test | FDA Guidelines | Minicircle Preparations |  |
| --- | --- | --- | --- |
|  |  | BMP2 | Luciferase |
| Residual host chromosomal DNA | The residual host chromosomal DNA content must be < 1.0 % (w/w) | < 1% | < 1% |
| Residual host RNA | The residual RNA content must be < 1.0 % (w/w) | < 1 % | 2 % |
| Residual host cells protein | The residual host cells protein content must be < 1.0 % (w/w). | below detection limit | below detection limit |
| **Purity test** |  |  |  |
| Plasmid topology | The supercoiled form percentage must be ≥ 80.0 %. | 97% | 92% and 96% |
| **Safety test** |  |  |  |
| Endotoxin content | The endotoxin content must be < 40 EU / mg of plasmid . | 0,16 EU/mg | 0,18 EU/mg |

The topology of the minicircle-DNA was investigated by HPLC as described in material and methods to estimate the proportion of supercoiled species after purification on DEAE and HIC columns. In case of the BMP2 minicircle preparation a fraction of 97% supercoiled DNA was measured by both methods, HPLC and CGE. In the case of the luciferase minicircle-DNA the same analysis were conducted resulting in an estimation of 96% and 92% supercoiled form as calculated by HPLC and CGE respectively. A chromatogram and an electropherogram of the purified luciferase minicircle-DNA in comparison with AGE analysis after digestion with the restriction enzyme HindIII can be seen in Figure 6. Quantification of residual host cell DNA by real time PCR revealed a gDNA content below the FDA maximum threshold of 1%. As specified by the FDA guidelines, the endotoxin content should not exceed 5 EU / mg of plasmid. Both, the BMP2 and Luciferase minicircle samples met these criteria as they only display a 0.16 and 0.18 EU/mg of plasmid respectively. The content of residual host cell protein was also below the FDA threshold of 1% (w/w) of the minicircle content. Both minicircle-DNA preparations met this requirement as the amount was below the detection limit of the ELISA method used for detection. Finally, analysis for RNA as described in material and methods revealed impurities below 1% (w/w) of the minicircle content in the case of the BMP2 minicircle-DNA preparation. Furthermore, analysis by AGE did not reveal RNA impurities in the luciferase minicircle-preparation shown in Figure 6a. The content of residual RNA must not exceed 1% (w/w) of the minicircle content. The minicircles derived from pRBPS-IVR7_BMP2 met that requirement, while the minicircle candidate derived from pRBPS-IVR7_ LucCmv contains up to 2% of residual RNA. These minicircles represent different products and are not prepared according to an optimized protocol for each construct as, according to the objective of this work, the development of a scalable protocol was the subject of research. Therefore, it is possible that the amount of impurities including the ratio between minicircle and residual RNA is not the same. For optimal final purity not only the final purification step, but also up-stream processes in sample preparation are crucial. Even though the content of residual RNA in the final luciferase minicircle sample was above the threshold, the case of the BMP2 minicircle preparation indicates that the required purity regarding RNA is certainly achievable with further optimisation of the steps in the process.

## DISCUSSION

The parental plasmid described in this work was developed to enable the stringent control of *in vivo* processes involved in minicircle-DNA production. The first process comprises site specific recombination of a parental plasmid to produce minicircle-DNA. The second process aims to remove the miniplasmid carrying the unwanted bacterial backbone sequences by enzymatic degradation. Due to the specific set-up of the parental plasmid described in this study, recombination and restriction are performed as coordinated, consecutive processes. A single induction step initiates a sequence of events, which finally result in a host cell harbouring almost exclusively minicircle molecules.

The problem that is addressed by this stringent process sequence is the premature degradation of the parental plasmid. Chen and coworkers were the first to publish an *in vivo* restriction approach to degrade the miniplasmid and residual (unrecombined) parental plasmid via co-expression of a site specific recombinase (PhiC31 integrase) and a homing endonuclease (I-SceI) (Chen et al. 2005). Both enzymes are located on the parental plasmid and are under the control of the P_BAD_ promoter of the arabinose operon. Thus, addition of the inducer, L-arabinose, results in the simultaneous expression of the integrase and the homing endonuclease. The problem of such a system set-up is that the parental plasmid, at least to some extent, is degraded by the endonuclease before recombination occurred thereby potentially reducing the yield of the desired product. To solve this issue, Chen et al took various measures to minimize premature degradation of the parental plasmid (Chen et al. 2005). First, an additional copy of the PhiC31 integrase gene was inserted into the plasmid system to accelerate the recombination process. Second, the cultivation temperature was increased from 32°C for site-specific recombination to 37°C for I-SecI endonuclease activity, as the recombination event is thought to be favoured at the lower temperature. Third, recombination was performed at a lower pH than the restriction reaction as I-SecI activity is known to be optimal at higher pH values (Monteilhet et al. 1990).

In a more recent approach an *E. coli* strain was modified to express the same set of inducible minicircle-assembly enzymes as mentioned above, namely the PhiC31 integrase and the I-SceI homing endonuclease (Kay et al. 2010). This producer strain carries ten copies of the integrase and three copies of the endonuclease. But even in this approach both enzymes are under the control of the same inducible PBAD/ araC arabinose promoter system described previously. Therefore, the same measures with regard to pH and temperature variation of the culture broth were necessary to minimize premature degradation of the parental plasmid (Kay et al. 2010). Furthermore, the stability of this genetically modified host strain might be an issue. Bacteria normally tend to loose redundant, unnecessary genomic information. The authors address this problem in their publication (Kay et al. 2010) by citing a study that shows that repeated gene sequences inserted into the bacterial genome are stable for 80 generations (Sallam et al. 2010). However, the cited study was performed in Rhodococcus using a different technique for chromosomal integration.

An alternative strategy for the coordination of recombination and restriction would be the use of different promoter systems to sequentially induce the expression of the involved enzymes. The main issue with this approach is the availability of appropriate promoter systems. Both enzymes need to be controlled very tightly as background expression leads either to premature recombination or to premature restriction, both resulting in complete loss of minicircle-DNA. Although there are a variety of inducible bacterial promoters known, only a few of them are suitable for such a purpose. Lactose promoter / operator based systems e. g. the lac, tac, trc and the T7 polymerase system (the latter is usually under the control of the lactose promoter) are known to be leaky and are therefore not suitable. Other systems like for example the tetA or the XylS promoter depend on inducer substances (tetracycline, m-toluic acid) which are problematic in terms of GMP production for clinical purposes, where only substances should be used that are generally regarded as safe.

The lambda promotor (λPR) is known for its stringent repression and is therefore a suitable candidate (Darquet et al. 1997; Nehlsen et al. 2006). However, the disadvantage associated with the lambda system is, that expression has to be induced by a temperature shift (usually from 37°C to 42°C) to inactivate the temperature sensitive mutant repressor used in this system. A temperature shift within a short time period is usually easy to perform with lab scale production volumes (e.g. 1 litre shaking flasks). However, a fast temperature shift as needed with such a system might constitute a significant effort in industrial large scale production, where bioreactors with volumes of several hundred and up to thousands of litres are operated.

A further point to consider in case two promoter/operator systems are involved is that the respective repressor molecules must be made available too. These molecules can be delivered in cis (i.e. by the same plasmid that contains the respective promoters) or in trans (i.e. by a second, compatible plasmid, by phage infection or via chromosomally integrated repressor expression cassettes). Both alternatives have disadvantages. Delivery via a second plasmid or via phages means that there are additional DNA-molecules involved which have to be removed in the course of minicircle-DNA purification. Chromosomal integration implies the disadvantage of limitation to (a) certain host strain(s). For delivery in cis, the expression cassette of both repressor proteins must be integrated into the plasmid carrying the two different promoter / operator systems. This would result in a very large parental plasmid encoding two repressors, the resolvase and the endonuclease, thus, the capacity for the insert (the gene of interest) would be very limited. Apart from this, timing of induction would be another point to consider. The optimal/right moment to induce endonuclease expression is obviously when recombination is completed. However, there is no monitoring process available that could indicate the endpoint of the recombination reaction during fermentation. Therefore, the question of the appropriate timing to induce the degradation process would hamper the usability of such a system. In the light of the so far discussed approaches, the parental plasmid presented in this report provides a reasonable solution to the problem of process coordination in the context of *in vivo* minicircle production.

With regard to fermentation we have identified optimal conditions for the production of two minicircle candidates using parental plasmids pRBPS-IVR7_BMP2 and pRBPS-IVR7_LucCMV at industrial pilot scale. In particular we showed that during the biomass production phase at 28°C both parental plasmids were stable and no degradation has been observed due to a tight arabinose promoter control preventing expression of the endonuclease which would have resulted in parental plasmid degradation. We have also shown for both plasmids that 3 hours at 37°C were sufficient to increase the plasmid number to a high copy number level. Additionally, we have demonstrated that both *in vivo* recombination and restriction worked very efficiently for both minicircle candidates upon induction with only 1 % L-arabinose although induction was performed at significantly elevated OD_600_ levels as compared to lab scale production. Interestingly, the results achieved with an induction period of 1.5 hours proved to be very similar as compared to 3 hours induction. Concerning product purification, the results achieved indicate that the developed process is scalable and enables also large scale production. It is especially worth mentioning that no miniplasmid has been detected in the analysed preparations demonstrating the efficiency of the degradation process. This is certainly a very encouraging result as the miniplasmid, due to restrictions in resolution, cannot be removed by means of conventional chromatography. Basically, following the purification steps both purified minicircle candidates met the ICH guideline except for luciferase minicircles which was displaying a slightly higher level of residual RNA in this particular preparation. Despite this fact, it was shown that monoliths play important role in purification of minicircle DNA. In this work monoliths were used as purification as well as analytical tool, the latter in combination with other well-known analytical techniques. The final samples produced by the purification process described herein represent products with high percentage of supercoiled DNA and with extremely low amounts of host cell DNA, endotoxins, residual host cell proteins and residual RNA.

It is already well established knowledge that minicircle-DNA is superior to conventional DNA, not only in terms of biological safety but also regarding transfection efficiency. In this study we used a reporter construct to demonstrate that minicircles produced with the RBPS *in vivo* restriction technology hold this promise. It should be pointed out that the minicircle-DNA showed significant higher luciferase expression levels at every measured time point when equimolar amounts (i.e. a considerably smaller total amount) where used. Although the same total amount (in μg) initially resulted in an even elevated luciferase activity as compared to the equimolar approach, this effect levels out after two weeks but on a much higher level as compared to the conventional plasmid. This can be interpreted as that with minicircle-DNA a much smaller amount of DNA is necessary and sufficient for gene transfer applications. This also means that with minicircle-DNA not only the burden of contaminating bacterial backbone DNA is reduced to a minimum but also the total amount of the necessary transfection agent is reduced, as compared to conventional pDNA. This seems important to notice as transfection reagents, as most of the delivery systems used in non-viral gene transfer, bear a toxic potential by themselves which can therefore be reduced as much less of these substances is needed in combination with minicircle-DNA.

With regard to the yield achieved in this study there is certainly some optimisation work left to do. With conventional plasmids, high production yields in the range of grams per litre have been reached in high cell density fermentations and are nowadays routinely achieved in the course of large scale production protocols. However, it must be mentioned that production yields of minicircle-DNA and conventional plasmid DNA cannot be easily compared as the production process of minicircles is far more complex. In contrast to conventional plasmids, minicircle-DNA production involves additional *in vivo* processes like site specific recombination and, as in the presented case, *in vivo* restriction and subsequent degradation of unwanted DNA molecules. Therefore, for minicircle production not only the maximum accumulation of plasmid DNA and high cell densities are important, but also the optimal condition for the host cell to produce the enzymes necessary and to drive the recombination and restriction/degradation processes. However, culture conditions necessary to achieve high quality recombination, restriction and finally degradation are not the same as for achieving high plasmid yields. Furthermore, minicircles are much smaller, usually a factor 2 – 3, a fact that needs to be adequately addressed in case total amounts are compared. And last but not least, as shown in this study a far smaller amount of minicircle DNA is necessary to show the same effect as with conventional plasmid DNA. Considering all these facts, the yields achieved in this pilot study are already in a very reasonable range indicating that the minicircle technology is able to replace conventional plasmids in clinical applications.

## ACKNOWLEDGEMENTS

We would like to thank Florence Xhonneux from Eurogentec for supporting this project as well as for critical review of the manuscript, Ara Hacobian from the Ludwig Boltzmann Institute for Experimental and Clinical Traumatology for kindly providing plasmid pEF1a-BMP2-advanced.

## COMPLIANCE WITH ETHICAL STANDARDS

### Research involving Human Participants and/or Animals

This article does not contain any studies with human participants or animals performed by any of the authors

